# Benchmarking DIA data analysis workflows

**DOI:** 10.1101/2023.06.02.543441

**Authors:** An Staes, Teresa Maia, Sara Dufour, Robbin Bouwmeester, Ralf Gabriels, Lennart Martens, Francis Impens, Simon Devos

**Affiliations:** VIB Center for Medical Biotechnology, Technologiepark-Zwijnaarde 75, B9052 Ghent, Belgium; Department of Biomolecular Medicine, Ghent University, Technologiepark-Zwijnaarde 75, B9052 Ghent, Belgium; VIB Proteomics Core, Ghent, Belgium

## Abstract

Data independent acquisition (DIA) has become a well-established method in LC-MS driven proteomics. Nonetheless, there are still a lot of possibilities at the data analysis level. By benchmarking different DIA analysis workflows using a ground-truth sample, consisting of a differential spike-in of UPS2 in a constant yeast background, we provide a roadmap for DIA data analysis of shotgun samples based on whether sensitivity, precision or accuracy is of the essence. Three different commonly used DIA software tools (DIA-NN, EncyclopeDIA and Spectronaut^TM^) were tested in both spectral library mode and spectral library-free mode. In spectral library mode we used the independent spectral library prediction tools PROSIT and MS2PIP together with DeepLC, next to the classical DDA-based spectral libraries. In total we benchmarked 12 DIA workflows. DIA-NN in library-free mode or using *in silico* predicted libraries, together with Spectronaut in library-free mode, shows the highest sensitivity maintaining a high reproducibility and accuracy. In general, DIA-NN shows the best reproducibility, while the accuracy is comparable for all DIA workflows.

## Introduction

Data-dependent acquisition (DDA) has been the golden standard for mass spectrometry driven proteomics for years. With an increasing number of samples needed for statistical analysis, especially for the analysis of large clinical cohorts, label-free quantification became the favored approach compared to label-based approaches. A challenge in label-free quantification is the high degree of missing values due to the stochastic nature of DDA^1, 2^. The repeatability in protein identification on Orbitrap instruments is only about 70-80%^1^ with DDA, as compared to 98-99% with data-independent acquisition (DIA)^3^. Tabb *et al.* showed that a minimum of three repeats of the same sample is needed to obtain sufficient information from an LC-MS/MS run in DDA^1^. One option to decrease the amount of missing values is through the use of isobaric labeling agents such as TMT^®^ or iTRAQ^®^. Although these reagents evolved in recent years (currently allowing the labeling of up to 18 samples in one run), the isobaric labeling approach suffers from ratio compression^4–6^. This problem can be tackled through extensive pre-fractionation or by running more sophisticated acquisition methods like synchronous precursor selection (SPS)^7^. Still these solutions either increase LC-MS/MS time tremendously or decrease the number of identifications, while being limited to a maximum number of 18 samples. When increasing the sample cohort above 18 samples^8^, different LC-MS/MS runs need to be done to compare the different TMT sets, inevitably running again into the stochastic nature problem of DDA and introducing more missing values.

A promising option to reduce the number of missing values without the use of isotopic labels is DIA. In DIA methods, all precursor ions from a certain m/z isolation window are simultaneously fragmented, and as such sampling the full m/z range and providing a solution for the stochastic nature of DDA. Indeed, studies demonstrated that the number of DIA identifications surpasses the theoretical maximal possible number of MS2 spectra that can be acquired on a Q-Exactive HF instrument with state-of-the-art DDA methods, using a tryptic Hela digest^3, 9^. Although very promising, the challenging analysis of DIA data hampered the widespread use of this acquisition method for a long time. By simplifying the data analysis to a targeted analysis using a DDA-created spectral library, DIA became more accessible^10, 11, 12^. However, this approach still needs upfront DDA analysis to create the spectral libraries, taking along its stochastic nature. Moreover, in order to generate deep ‘project-specific libraries’ a lot of sample material and MS time is consumed^3^.

The use of publically available ‘resource spectral libraries’ reduces sample consumption and analysis time, and can lead to a 90-103% performance of protein identification as compared to project-specific libraries^3^. On the other hand, quantification is still more precise when using project-specific libraries^3^. Also, the resource spectral libraries are generated under variable conditions with variable quality, and are thus prone to large false-discovery rates (FDR)^13–15^. In recent years, the creation of spectral libraries through machine learning gained much attention. When creating a spectral library *in silico*, a complete database can be covered that is applicable for DIA analysis, while uncoupling it from upfront DDA analysis and its restrictions. Algorithms such as MS2PIP^16, 17^, PeptideArt^18^, DeepDIA^19^, DeepMass:PRISM ^20^ and PROSIT^21^ all use (deep) machine learning to create *in silico* spectral libraries. These libraries can also be merged with project/resource spectral libraries to generate hybrid spectral libraries. Nowadays, some data analysis software even have an embedded ‘library-free’ search option where peptides are identified directly from the DIA data. When using predicted spectral libraries, the success rate depends on how well the predicted spectra match the actual spectra. Considering the large amount of precursors that are present in such a database, the FDR needs to be controlled more extensively then^22^. DIA data analysis can be further improved through the prediction of peptide retention times^20, 23, 24^ with algorithms like Elude^25, 26^ and DeepLC^27^, or by using chromatogram libaries^24, 28^.

Next to the way the spectral library is created, the DIA software approaches can be divided in two main categories in the way it performs the analysis: a peptide-centric or a spectrum-centric approach. A peptide-centric approach will look for a peptide across all isolation windows through the alignment of the retention times of the fragment ions readily available in the spectral library. Concurrent elution profiles point to the same peptide. A spectral library is required to perform this targeted approach. Examples of software tools using this approach are PECAN (or its improved version Walnut, embedded in EncyclopeDIA)^18, 23, 29^, Skyline^10, 30^, MSPLIT-DIA^31^, OpenSwath^12^ and MaxDIA^20^. In contrast, a spectrum-centric approach will look for peptide and fragment ion features in a particular isolation window, leading to the selection of the single best matching precursor for each spectrum, called pseudo-MS2 spectra^31, 32^. These pseudo-MS2 spectra can then be searched with DDA search algorithms. Software using a spectrum-centric approach are DirectDIA (in Spectronaut), DIA-Umpire^33, 34^ and Group-DIA^35^. DIA-NN^32^ uses both the peptide-centric and spectrum-centric approach, combining the best of both worlds. Furthermore, DIA-NN predicts *in silico* spectral library from a fasta file, as is also the case for MaxDIA^20^. Although DIA-MS largely focuses on the MS2 level, the MS1 information can be used as well for quantification purposes and for filtering out possible interferences. While Spectronaut makes use of this former approach on the MS1 level only, DIA-NN uses an interference filter on both the MS1 and MS2 level^24, 32, 36^. EncyclopeDIA uses chromatogram libraries, using the Peptide Query Parameters (PQP) information to improve the quality of the DIA analysis.^23, 24, 37^. Since retention time information was proven to be powerful in chromatogram libraries^23, 24^, and in DIA analysis in general ^20^, the *in silico* spectral library creation tools MS2PIP and PROSIT both have an embedded retention time prediction tool included to add retention time information to the *in silico* created spectral libraries. Now that DIA data analysis becomes much more powerful, it has proven its success in the analysis of clinical samples ^38–43^ and shows great promise in single cell proteomics^44, 45^.

In order to create better insights into the advantages and disadvantages of the different DIA data analysis workflows, we evaluated different approaches that are currently available for DIA data analysis in terms of detectability, precision and accuracy. The use of independent, *in silico* generated spectral libraries was also included here, which was not reported before. For this, a suitable benchmark sample was created that mimics a real proteome sample. The Universal Protein Standard 2 (UPS2), containing 48 human proteins with a dynamic range spanning six orders of magnitude, was spiked in a constant yeast background in different amounts, and was measured with both DDA-MS and DIA-MS. This to compare the classical DDA data analysis workflow with 12 different DIA data analysis workflows. The classical DDA data analysis was performed with MaxQuant, while the DIA data analysis workflows was performed with the commonly used software tools Spectronaut^TM^, DIA-NN and EncyclopeDIA. All three tools were run with four different spectral libraries modes: a DDA-library mode, a tool-specific library-free mode, an MS2PIP-generated *in silico* spectral library mode and an PROSIT-generated *in silico* spectral library mode. By analyzing dilution series of the spiked UPS2 standard we mimic true biological proteome samples, and therefore get a more realistic view on the different performance parameters of the DIA workflows and their limitations, as compared to other benchmark studies^11, 46, 47^.

## Material and methods

### Sample preparation

One vial of the Proteomics Dynamic Range Standard Set (UPS2; Sigma), containing 6 groups of 8 proteins in an amount of either 50000 fmol, 5000 fmol, 500 fmol, 50 fmol, 5 fmol and 0.5 fmol (UPS2 abundance groups; Table 1), was dissolved in 200 µl 50 mM triethylammonium bicarbonate (TEAB) (Sigma), after which 400 ng of trypsin (Promega) was added. After incubating the sample overnight at 37°C, digestion was stopped by adding 10 µl of 10 % trifluoroacetic acid (TFA). The peptide mixture was further diluted to a final volume of 400 µl with LC-MS loading buffer (0.1% TFA in 2:98 (v:v) acetonitrile (ACN)/water). The digested UPS2 sample was then spiked at different concentrations in a yeast digest background (Sigma), each time by dissolving 2 µg of dried yeast tryptic digest in 80 µl of the different UPS2 dilutions (1x, 1.25x, 2.5x, 5x and 10x dilution were prepared in LC-MS loading buffer; Table 1). Finally, 4 µl of indexed retention times (iRT) peptides was added to each sample of the UPS2 dilution serie (Biognosys; diluted according to manufacturer protocol).

**Table 1.** Per-protein amount of digested UPS2 sample analyzed with LC-MS, present in the different abundance groups and dilutions.

| Dilution factor | UPS2 abundance group |  |  |  |  |  |
| --- | --- | --- | --- | --- | --- | --- |
|  | 50000 | 5000 | 500 | 50 | 5 | 0.5 |
| 1x | 1.19 pmol | 119 fmol | 11.9 fmol | 1.19 fmol | 119 amol | 11.9 amol |
| 1.25x | 0.95 pmol | 95 fmol | 9.5 fmol | 0.95 fmol | 95 amol | 9.5 amol |
| 2.5x | 0.48 pmol | 48 fmol | 4.8 fmol | 0.48 fmol | 48 amol | 4.8 amol |
| 5x | 0.24 pmol | 24 fmol | 2.4 fmol | 0.24 fmol | 24 amol | 2.4 amol |
| 10x | 0.12 pmol | 12 fmol | 1.2 fmol | 0.12 fmol | 12 amol | 1.2 amol |

The Universal Proteomics Standard Set 1 (UPS1; Sigma) was digested in the same way as UPS2. To 40 µl UPS1 digest, 2µl of iRTs was added.

### LC-MS analysis

Each sample (5 µl of UPS1 or UPS2 digest, 10 µl of UPS2-spiked yeast digest) was injected on an Ultimate^TM^ 3000 RSLC ProFlow nano-LC system in-line connected to a Q Exactive^TM^ HF BioPharma mass spectrometer (Thermo). First, peptides were trapped for 4 min on an in-house packed C18 ReproSil-HD trapping column (5 µm, 100Å, 0.1x2 mm IDxL; Dr. Maisch, Germany) using loading solvent A (0.1% TFA in ACN/water (2:98, v/v) at 10 µL/min. After trapping, the peptides were loaded and separated on an analytical, in-house packed C18 ReproSil-Pur analytical column (1.9 µm, 100Å, 0.075x400 mm IDxL; Dr. Maisch, Germany), equipped with a laser-pulled electrospray tip using a P-2000 Laser Based Micropipette Puller (Sutter Instruments).The column was kept at a constant temperature of 50°C. Peptide separation was established through a stepwise gradient using solvent A (0.1% FA in water) and solvent B (0.1% formic acid (FA) in water/acetonitrile (2:8, v/v): from 0-30% solvent B in 105 min, from 30-56% solvent B in 40 min and from 56-97% solvent B in 5 min, followed by a 10-min wash with 97% solvent B and re-equilibration with solvent A. During the whole gradient the flow rate is set at 250 nL/min. For both analyses, a pneu-Nimbus dual column ionization source was used (Phoenix S&T), at a spray voltage of 3.5 kV and a capillary temperature of 275°C. Both for DDA and DIA, the same sample amount was injected, and the same LC and ESI parameters were applied.

### DDA-MS

Full-scan MS spectra (375-1500 *m*/*z*) were acquired at a precursor resolution of 60,000 at 200 *m*/*z* in the Orbitrap analyser after accumulation to a target value of 3E6 with a maximum ion time of 60 ms. The 16 most intense ions above a threshold value of 13,000 were isolated with a 1.5 m/z window for higher-energy collisional dissociation (HCD) fragmentation at a normalized collision energy (NCE) of 30% after accumulating ions at a target value of 1E5 for a maximum of 80 ms injection time using a dynamic exclusion of 12 s. MS2 spectra (200-2000 *m*/*z*) were acquired at a resolution of 15,000 at 200 *m*/*z* in the Orbitrap analyser.

### DIA-MS

Full-scan MS spectra (375-1500 *m*/*z*) were acquired at a precursor resolution of 60,000 at 200 *m*/*z* in the Orbitrap analyser after accumulation to a target value of 5E6 with a maximum ion time of 50 ms. After each MS1 acquisition, thirty 10-m/z isolation windows (with an overlap of 5 m/z) were sequentially selected by the quadrupole for HCD fragmentation at a NCE of 30% after filling the trap at a target value of 3E6 for a maximum injection time of 45 ms. MS2 spectra were acquired at a resolution of 15,000 at 200 *m*/*z* in the Orbitrap analyser without multiplexing. The isolation windows were acquired over a mass range of 400-900 m/z (covering over 90% of a tryptic digest^10, 24, 48^). The isolation windows were created with the Skyline software tool (v3.6).

### DDA data analysis

DDA-MS data was analysed with MaxQuant (version 2.2.0.0) using mainly default search settings, including a FDR set at 1% on PSM and protein level. Spectra were searched against a *Saccharomyces cerevisiae* protein database (UniProt database release version of January 2023 containing 6,727 yeast protein sequences, downloaded from http://www.uniprot.org; hereinafter referred to as “*Saccharomyces cerevisiae* protein sequence database*”*), and the protein sequences of the 48 human proteins from UPS2 (provided by Sigma; hereinafter referred to as “UPS protein sequence database*”*). The mass tolerance for precursor and fragment ions was set to 4.5 and 20 ppm, respectively, during the main search. Enzyme specificity was set as C-terminal to arginine and lysine, also allowing cleavage at proline bonds, with a maximum of two missed cleavages allowed. Variable modifications were set to oxidation of methionine residues, acetylation of protein N-termini and carbamidomethylation of cysteines. For DDA data analysis, ‘matching between runs’ was enabled, with a matching time window of 0.7 minutes and an alignment time window of 20 minutes. MaxLFQ was enabled as well, with a minimum ratio count of two unique or razor peptides required for quantification.

### DDA-based spectral library creation

In MaxQuant, a DDA spectral library was created for DIA using the same search parameters as described above, except for the ‘matching between runs’ and ‘MaxLFQ’ options, which were disabled. In order to create a more comprehensive DDA spectral library, both the UPS2 sample (dynamic range standard) as well as the UPS1 sample (equimolar standard) were included in the searches.

In the case of Spectronaut (version 17.2), the DDA based library was created with the embedded tool using default parameters. The library was created through the *msms* file of the dedicated MaxQuant search. For EncyclopeDIA (v1.2.2), a dlib DDA-based library was created from an *msms* file and a *fasta* file containing all *Saccharomyces cerevisiae* and UPS protein sequences, using default settings. A further precursor mass filter of 400-900 m/z was applied to the spectral library file, using the DB Browser for SQLite software (https://sqlitebrowser.org/49). In DIA-NN the *msms* file was used as well, embedded in the search to create a DIA-NN-compatible DDA-based library.

Table 2 summarizes the number of precursors present in each database.

**Table 2.** Number of unique precursors in each library for the different workflows in the DIA analysis.

| DIA workflow | Number of precursors |
| --- | --- |
| DIA-NN DDA library | 30,926 |
| DIA-NN MS2PIP library | 1,500,882 |
| DIA-NN PROSIT library | 648,619 |
| DIA-NN library-free | 969,682 |
| Spectronaut DDA library | 3019 |
| Spectronaut MS2PIP library | 806,672 |
| Spectronaut PROSIT library | 646,453 |
| Spectronaut library-free | 3,822 |
| EncyclopeDIA DDA library | 25,383 |
| EncyclopeDIA MS2PIP library | 793,314 |
| EncyclopeDIA PROSIT library | 648,619 |
| EncyclopeDIA library-free | 201,998 |

### *In silico* predicted spectral library creation

#### MS2PIP

From both the UPS protein sequence database and the *Saccharomyces cerevisiae* protein sequence database, an *in silico* spectral library was created using the MS2PIP algorithm^50, 51^ (v3.11.0) in combination with the DeepLC algorithm (DeepLC v1.2.1, DeepLC Retrainer v0.1.13, and Pyteomics 4.5.6) using the retention time of the identified peptides from all DDA runs. For EncyclopeDIA (v1.2.2), a dlib library was then created on the software GUI from the msp format. A further precursor mass filter of 400-900 m/z was applied to the spectral library file, using the DB Browser for SQLite software (https://sqlitebrowser.org/49).

#### Prosit

The Prosit^52^ *in silico* library was generated in two steps. First, the UPS and the *Saccharomyces cerevisiae* protein sequence databases were used to create a Prosit CSV file in the DIA-NN software, with ‘Default NCE’ set to 34 (optimal collision energy value defined by the CE calibration tool from the Prosit website (https://www.proteomicsdb.org/prosit/), based on search results from one of the DDA runs) and ‘Maximum Missed Cleavages’ set to 2. Next a library was obtained from the Prosit website, using the Prosit_2020_intensity and Prosit_2019_irt prediction models, and a PROSIT-predicted NCE of 34. This was exported in comma delimited, Spectronaut-compatible input format.

#### Use of in silico predicted libraries

In Spectronaut (v17.2) the created *in silico* predicted spectral library was imported with the embedded spectral library tool using default settings, except for the “proteotypicity filter” which was checked. The extra option ‘*in silico library*’ was checked upon searching. The application of the *in silico* predicted spectral libraries in DIA-NN (v1.8.1) required the msp format in the case of MS2PIP after concatenation of the UPS and yeast msp files. The reannotation option was used with the UPS and *Saccharomyces cerevisiae* protein sequence databases. In the case of PROSIT, the csv file was first reannotated in DIA-NN creating a tsv format spectral library. In EncyclopeDIA (v1.2.2) the dlib format, created as described above, was used as spectral library.

### DIA data analysis

Thermo experimental RAW files were converted in mzML format using msconvert^53^ from the Proteowizard software package^54^, with the following parameters: binary encoding precision set at 32, SIM as spectra, peak picking using vendor libraries at MS 1 level, and demultiplex overlapping spectra. The resulting mzml files were used for analysis in DIA-NN and EncyclopeDIA. In Spectronaut the files were demultiplexed through the software starting from the RAW files (details see further). Table 2 summarizes the number of precursors that were used as target in the different workflows.

#### DIA-NN

DIA-NN (v1.8.1) was used through the graphical user interface. In case of the library free search, the FASTA digest option was checked with the UPS protein sequence database and the *Saccharomyces cerevisiae* protein sequence database added. For all workflows, trypsin was selected as the digestion enzyme (allowing cleavage N-terminally at proline and allowing up to two missed cleavages), and carbamidomethylation of cysteines, oxidation of methionines and acetylation of N-termini were set as variable modifications. Mainly default search settings were used, except the precursor mass range filter which was set at 400-900 m/z, the match between runs (MBR) option which was enabled, and “mass accuracy“ (MS2 mass accuracy) which was set at 20 ppm and “MS1 accuracy“ (MS1 mass accuracy) at 10 ppm. Only proteins identified with 2 peptides and with a protein group q-value below 0.01 (“Lib.PG.Q.Value”) were retained in the report for further analysis using the MaxLFQ value of each remaining protein group.

#### Spectronaut

For the DIA analysis in Spectronaut (v17.2), experiment files were first converted from the mass spectrometer vendor’s raw format to the HTRMS format through the HTRMS converter (v17.2), using default parameters with MS2 demultiplexing enabled. The DIA analysis was done with default settings, except for the protein LFQ method which was set to MaxLFQ in order to maximize quantification reproducibility^55, 56^. Only in the case of *in silico* generated libraries, the *in silico* optimization parameter was checked. For the DirectDIA (v17.2) analysis the vendor raw format was used. The analysis was done with default parameters, except for MS2 demultiplexing which was checked, in addition to the proteotypicity and fasta matched as a quantification prerequisite. The fasta files used for each analysis workflow contained the UPS protein sequence database and the *Saccharomyces cerevisiae* protein sequence database. The following variable modifications were set: acetylation on protein N-termini, oxidation of methionine, and carbamidomethylation of cysteines. The default pivoted protein report was used as report. Values used for further analysis were “PG. Quantity”.

#### EncyclopeDIA

EncyclopeDIA (v1.2.2) was run through the command line, using mostly default settings, except for a precursor and a fragment mass tolerance of 15 ppm and 20 ppm, respectively. The UPS and the *Saccharomyces cerevisiae* protein sequence fasta databases were specified. For the library-free search the ‘walnut’ parameter was enabled, while for the library-dependent searches the path to the corresponding dlib spectral library file was set. Further analysis was done with median normalized protein quantification values per protein group.

## Results

### Experimental setup

To compare classical DDA analysis with different DIA workflows, a benchmark sample was used containing the Universal Protein Standard 2 (UPS2) standard spiked in a constant yeast background in different amounts. The UPS2 sample was digested with trypsin and spiked in a commercial yeast tryptic digest at five different ratios ranging from 1:10 to 1:1. Next to this different spike-ins, the 48 human proteins in UPS2 are already spanning six orders of magnitude (6 groups of 8 proteins). This results in both inter- and intra-differential abundancies of the 48 proteins in the different samples ranging from 1.2 amol to 1.2 pmol. The dilution series covers a larger dynamic range as compared to Gotti *et al.*^46, 57^, consequently providing a more realistic view on the strengths and limits of the different workflows. Moreover, such sample resembles a real proteome sample containing differential abundant proteins more closely as compared to the triple proteome LFQ benchmark sample, used for other DIA benchmarks^11, 47^. Indeed, in a typical differential shotgun analysis experiment, the sample proteomes are not regulated to a large extent, except for a handful of proteins. Hence, because of a ground-truth set of only 48 proteins at different concentrations, a better estimation of the limits of detectability, accuracy and precision is obtained.

### Contemporary DIA data analysis workflows

Currently, two different DIA data analysis workflows are commonly used: DDA-based spectral library analysis and spectral library-free analysis. To be able to test predicted spectral libraries we used two algorithms capable of generating *in silico* generated spectral libraries, MS^2^PIP and PROSIT. Each of these spectral library workflows can be performed with several algorithms available for DIA data analysis. Combining the possibilities both on the software level and on the workflow level results in many data analysis options and constitutes an important limitation for the wider implementation of DIA. In order to get a better view on the advantages and disadvantages of the different options, the performance of 12 different DIA data analysis workflows was evaluated, together with a classical DDA workflow, on the benchmark sample described above (Figure 1). The 12 DIA workflows consist of four data analysis workflows (DDA library, MS2PIP-predicted *in silico* library, PROSIT-predicted *in silico* library, library-free), executed by three widely used software packages: Spectronaut^TM^ ^58^, EncyclopeDIA^28^ and DIA-NN^59^. Spectronaut^TM^ is a commercial software package using a spectrum-centric approach, while EncyclopeDIA is freeware using a peptide-centric approach. DIA-NN is freely available as well and combines both peptide- and spectrum-centric approaches.

**Figure 1.**
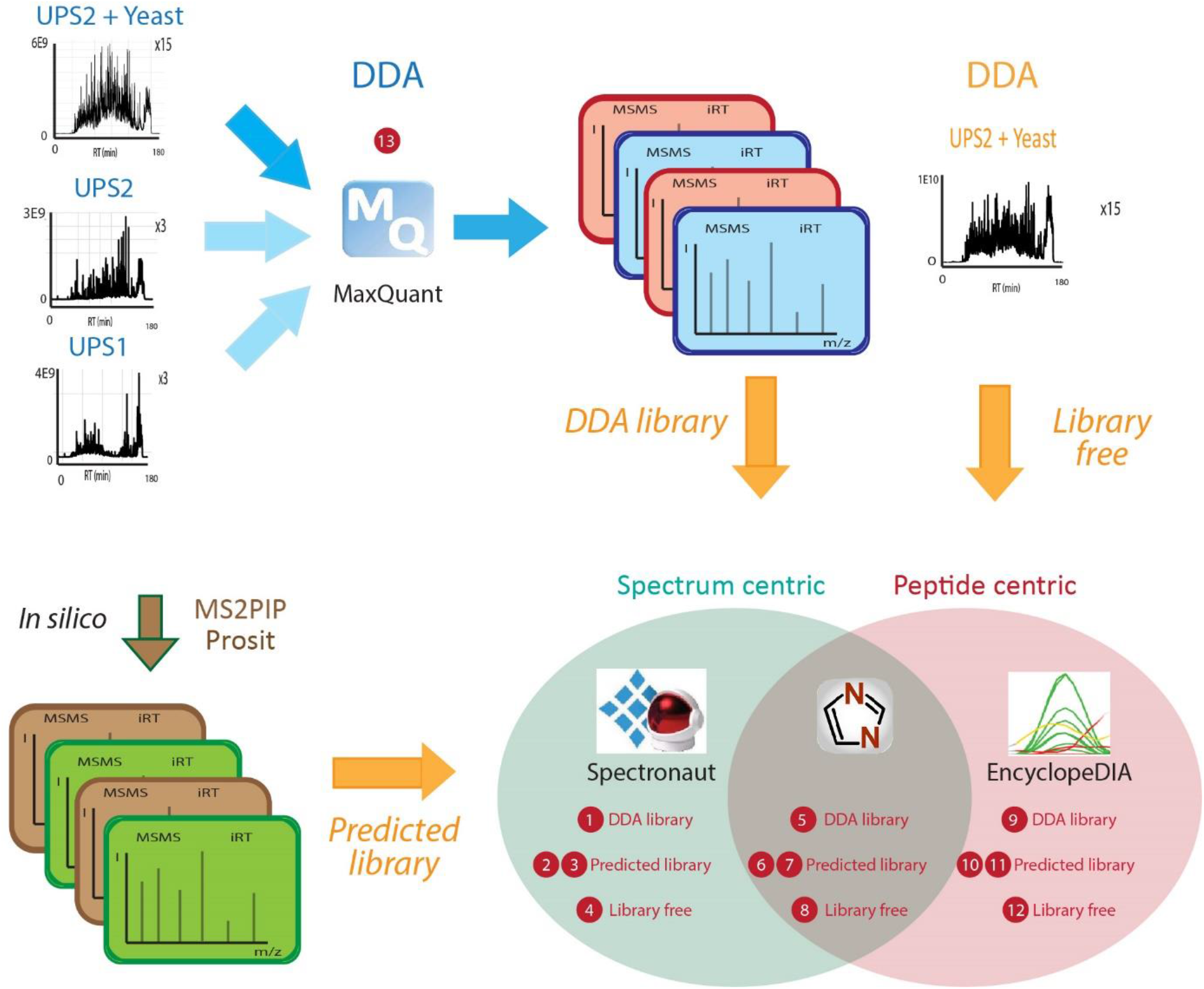
Experimental setup. DIA benchmark samples were tryptic digests composed of a standard of 48 recombinant proteins ranging six orders of magnitude (UPS2 reagent, Sigma), spiked into a *Saccharomyces cerevisiae* protein digest background. Five different samples were used, with the following dilution factors of UPS2 mixture in yeast extract: 1x, 1.25x, 2.5x, 5x and 10x. The DDA analysis methodology was compared side by side with a total of 12 DIA analysis workflows, differing on the type of spectral library option used as well as on the software. Spectral library options were: 1) DDA-based spectral library prepared with MaxQuant, 2) No spectral library/library-free, 3) MS2PIP-predicted spectral library, 4) PROSIT-predicted spectral library. In turn, software was MaxQuant for DDA data, while DIA data was analyzed on Spectronaut Pulsar, EncyclopeDIA or DiaNN. Both DDA and DIA runs were performed in triplicate on a Q Exactive HF mass spectrometer. For the DDA library creation, UPS1 and UPS2 were analyzed in DDA separately in triplicate as well.

On one hand, the benchmark sample was acquired and analyzed in triplicate with a DDA workflow, and on the other hand acquired in DIA and analyzed with each of the 12 different DIA workflows. The DDA data was analyzed with MaxQuant and served as the base for the DDA-derived spectral library construction. The DDA spectral library contained not only data from the DDA-MS analysis of the UPS2 sample (with and without yeast background), but from the UPS1 sample as well (without background). The UPS1 sample contains the same 48 human proteins but in equimolar amounts, which increases the depth of the library. Here, the chance that all 48 proteins are detected and identified in the DDA run is higher, and hence results in a more comprehensive DDA spectral library. The addition of the two separate DDA runs also mimics spectral library creation from a fractionated sample, a frequently applied approach^3^. However, to compare the performance of the DDA workflow to the different DIA workflows, the separate UPS1 and UPS2 analysis were excluded. *In silico* libraries created outside all DIA software were created with two different established algorithms: Prosit^52^ and MS2PIP^50, 51^ (the latter with the embedded DeepLC^27^ prediction tool to cover the retention time prediction).

Each workflow was both qualitatively (detectability) and quantitatively (precision and accuracy) evaluated using the differentially spiked UPS2 protein set. In addition, the (constant) yeast proteome background was assessed as well since it comprises 15 replicates in total.

### Detectability

The detectability of the different workflows was evaluated by the number of quantified human proteins of the set of 48 UPS proteins, identified with at least 2 peptides (Figure 2). A clear drop in the number of UPS2 protein identifications is noticed for the DDA workflow at the lowest spiked concentration, whereas for the other spike-ins the number is consistent. For all DIA workflows, the number of quantified proteins is higher as compared to the DDA workflow. The number of quantified UPS proteins in DIA is however more easily influenced by the spike-in dilution, except for the library-free mode in Spectronaut and EncyclopeDIA, which are more stable and hardly showing any dilution effect regardless of the analysis mode that is used. All DIA-NN workflows show decreasing numbers of quantified UPS proteins from high to low spike-in amount. Spectronaut identifies equal amounts of proteins for all spike-in amounts in the library-free approach, and shows decreasing numbers according to the spike-ins for all other workflows. DIA-NN outperforms all other approaches in the highest spike-in, except when using the DDA library, which performs best in Spectronaut. On the lowest spike-in amounts, Spectronaut performs best in DDA library mode and library-free mode, while DIA-NN and EncyclopeDIA performs best when using *in silico* created libraries.

**Figure 2.**
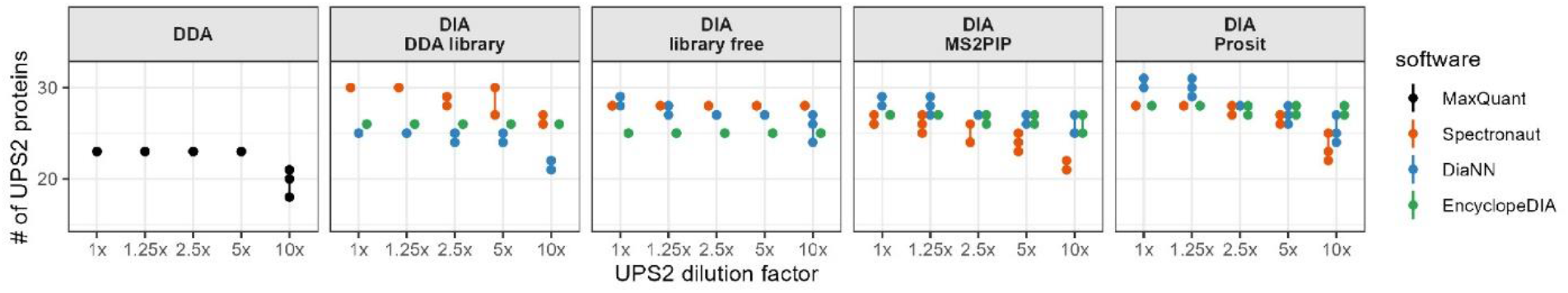
Detectability of the UPS2 proteins according to analysis workflow and sample dilution. Total number of UPS2 standard proteins (n=48) identified in each analysis workflow, according to sample dilution in yeast extract (1x down to 10x). Each software is represented with a different color. Panel columns depict the DDA workflow and the three DIA library methods.

In order to have a better view on the detectability of the lower abundant proteins, the data was separated according to the abundance of the UPS proteins (abundance groups 50000, 5000, 500, 50, 5, 0.5; Supplementary Figure 1). As expected, the lower abundant proteins are more prone to remain undetected. This analysis confirms that DDA suffers more from missing values than DIA in general, especially on low abundant proteins. The DDA signal starts to drop around 1.2 fmol and even disappears at 120 amol depending on the set of proteins. Still, DIA has its own limitations and is highly dependent on the algorithm and workflow that is used. With DIA-NN, the library-free and the two *in silico* predicted library approaches have the highest sensitivity, which is 9.6 amol in this setup. Spectronaut detects the most sensitive when using the PROSIT-generated *in silico* library and using a DDA library, going as low as 9.6 amol. In Spectronaut, the MS2PIP-created library approach and library-free approach does not detect anything down from the second last abundant set of proteins. EncyclopeDIA detects the second lowest set of proteins equally well in all spike-in amounts when using a DDA library or a PROSIT-predicted *in silico* library, but in library-free mode and using the MS2PIP-generated library this set of proteins is not detected anymore. The difference in performance in the DIA workflows is most visible from the third lowest abundant set of proteins onward. On this abundance set, Spectronaut outperforms the other software tools when using the library-free mode, while for all other modes DIA-NN shows the best performance. In general, despite the high number of identifications, still not all 48 UPS proteins are quantified in any workflow.

To summarize, DIA-NN with *in silico* predicted libraries and in library-free mode shows the highest sensitivity for only one UPS protein at an amount of 9.6 amol for two replicates. Both EncyclopeDIA and Spectronaut are able to detect proteins at an amount of 12 amol with the DDA-library and with the PROSIT-generated *in silico* library.

### Precision

The precision of MS2 quantification is reported to be higher as compared to MS1 quantification^36^. Therefore, besides evaluating the detectability in DIA versus DDA, the reproducibility of the quantification results was also evaluated. For each protein, a normalized intensity CV was calculated, per sample and per workflow (limited to the presence of three quantification values). Figure 3 shows the boxplots of the CVs for all commonly identified UPS proteins. The UPS2 protein set was also separated according to the abundance of the UPS proteins (abundance groups 50000, 5000, 500, 50, 5, 0.5) and the CV values of each group were represented (Supplementary Figure 2). All workflows manage to obtain CV values below 20%. In general, DIA-NN has the lowest CVs. remaining below 10%, and in most cases outperforms DDA itself. The CV seems to be independent of the library mode used for DIA-NN. EncyclopeDIA shows the highest CV values for all approaches, but remains below 20% in most cases. EncyclopeDIA as well seems to have CV values that are independent of the library mode used. The reproducibility for Spectronaut is more depending on the workflow, with the library-free mode showing the best precision for this software tool. Considering the different abundancies of the sets of UPS proteins, it shows that the CV values increase with decreasing protein abundancy. For the lowest abundance set of proteins for which a CV can be determined, DIA-NN is still outperforming the two other software tools.

**Figure 3.**
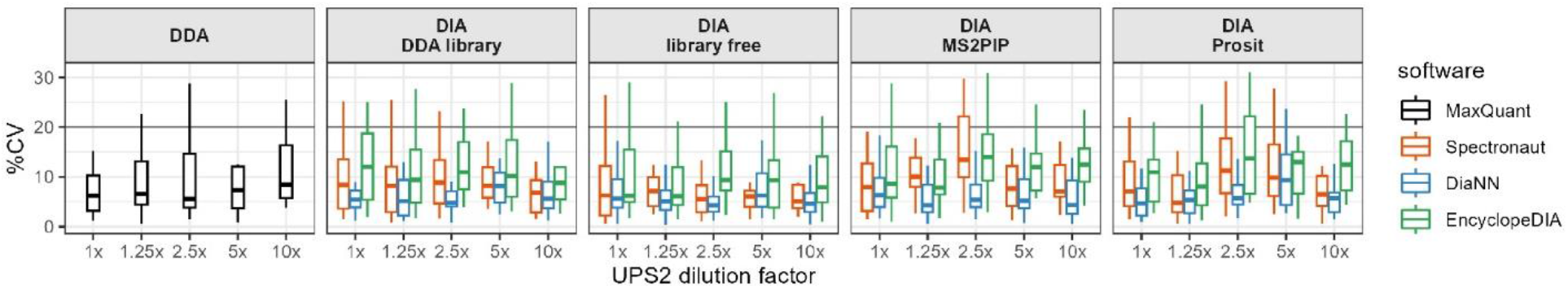
Quantification precision of the UPS2 proteins according to analysis workflow and sample dilution. Boxplots of percent coefficients of variation (%CV) of UPS2 mixture proteins (n=48) identified in each analysis workflow, according to the dilution in yeast extract (1x down to 10x). Plots’ horizontal lines are at 20 %CV.

All workflows suffer from higher CVs towards the lower spike-in concentrations. Indeed, for the lower abundant proteins a less intense signal is expected, which in turn leads to higher CV values. In the abundance set where DDA starts to suffer the most from lower detectability (the third lowest abundancy set) DIA-NN workflows remain to have a CV below 20% except for one spike-in (the 5x). The lowest abundant protein detected still has a CV below 20%. We can conclude that the extra identifications in DIA workflows still come with an acceptable precision, especially when using DIA-NN.

### Accuracy

Because the UPS2 set was spiked in the yeast background at different concentrations, it creates the possibility to examine the accuracy of inter sample protein quantification across the dilution series. The Mean Absolute Percentage Error (MAPE) was used to create an estimate of the error on the ratio of the intensities of the same UPS proteins between samples, using the undiluted (1x) sample as a reference.

MAPE values were combined in boxplots for all commonly identified UPS proteins (Figure 4). The UPS2 protein set was again separated according to the abundance groups and the MAPE values of each group were represented in boxplots (Supplementary Figure 3). DDA suffers less from the higher dilutions as compared to DIA. This is probably mainly due to the higher sensitivity of DIA in general. Indeed, low abundant proteins will suffer from a lower S/N ratio, to a large extent interfering with the area determination of the low intensity signals. Spectronaut has the lowest accuracy on the UPS ratios, especially with the DDA library mode. EncyclopeDIA and DIA-NN perform equally well in terms of accuracy in all modes, having a decreased accuracy with the lower spike-ins. Higher spike-ins of the low abundant UPS proteins still show a good accuracy.

**Figure 4.**
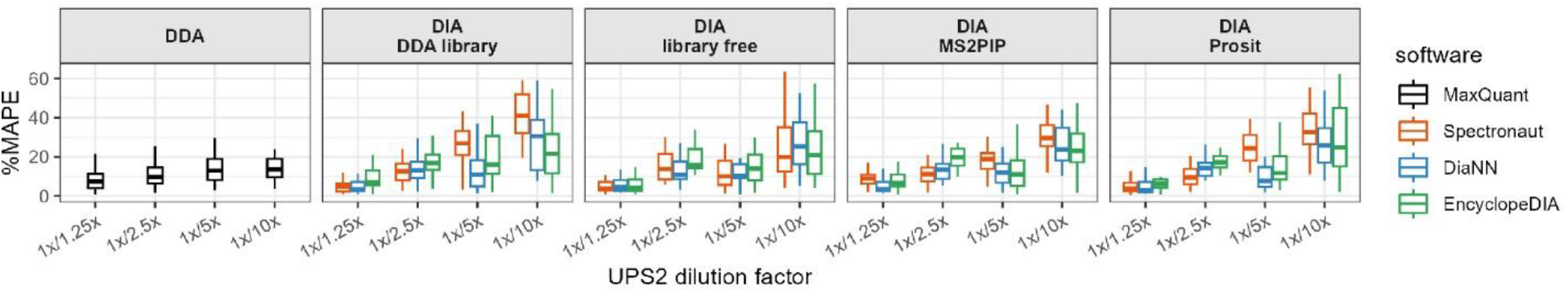
Quantification accuracy of the UPS2 proteins according to analysis workflow and sample’s protein abundance. Assessment of protein group quantification accuracy was done using the median absolute percentage error (MAPE) of the ratios of UPS2 protein intensities between the 1x yeast extract:UPS2 dilution relative to each of the other four sample dilutions (1.25x, 2.5x, 5x, 10x).

### Evaluation of yeast background

Next to the results on the UPS proteins, we have 15 replicates (5 dilutions, 3 replicates) of the yeast data as well (Figure 5). Although the accuracy cannot be determined, the detectability and precision can be evaluated. DIA-NN in library-free mode and using *in silico* predicted libraries outperforms all other DIA workflows by far in terms of detectability (Figure 5A). The use of the *in silico* predicted libraries renders higher protein numbers in general, but only DIA-NN has a comparable performance when comparing *in silico* predicted library mode to library-free mode. DDA library mode renders the lowest number of identifications in the DIA workflows, with EncyclopeDIA being the best to cope with a DDA library. DDA is outperformed by all DIA workflows, except for Spectronaut using the DDA library and using library-free mode.

**Figure 5.**
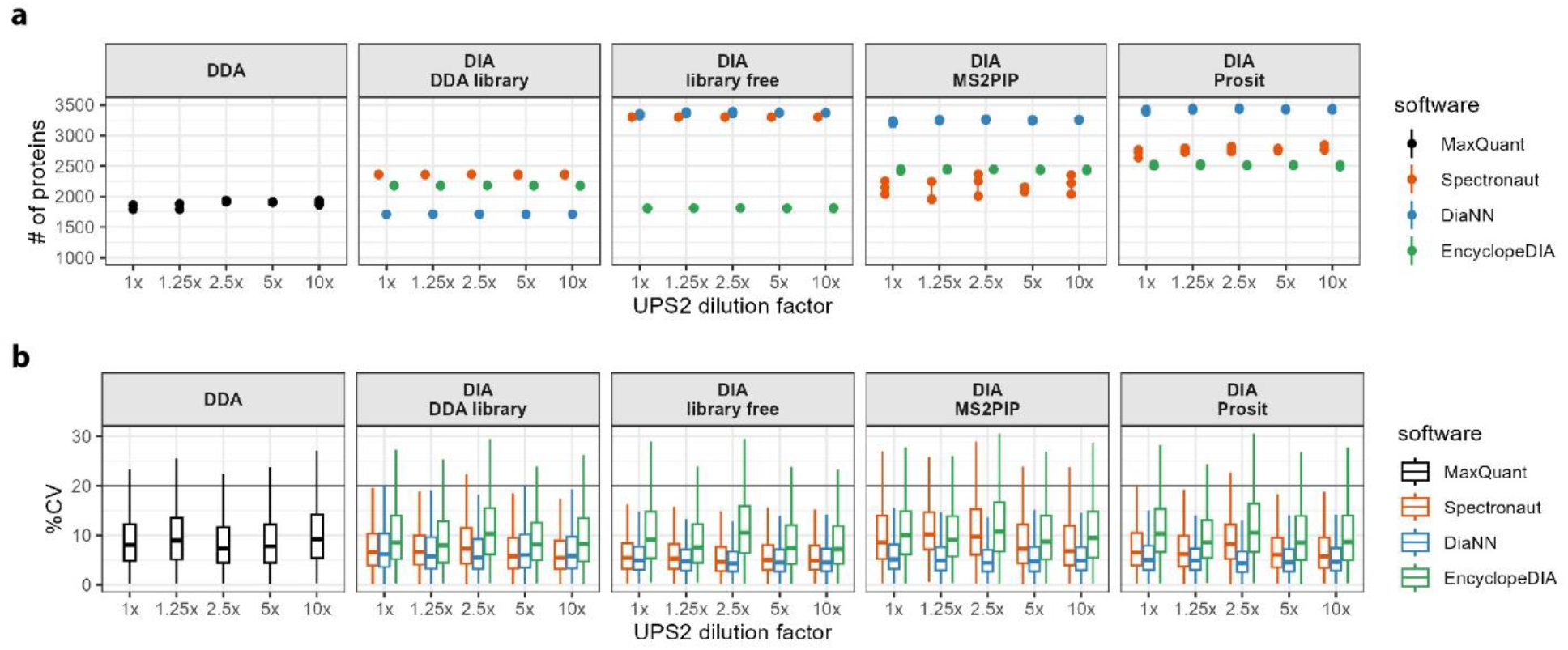
Detectability and quantification precision of background yeast protein groups according to analysis workflow. a. Total number of identified proteins in each analysis workflow, according to sample type (10x down to 1x dilution of UPS2 mixture in yeast extract). Each software is represented in a different color. Each column depicts a different library method and the DDA workflow. b. Boxplots of percent coefficients of variation of yeast proteins identified in each analysis workflow, according to sample type (10x down to 1x dilution of UPS2 mixture in yeast extract).

On the precision aspect (Figure 5B), DIA-NN outperforms all other workflows, including the DDA workflow. Encyclopedia is comparable in DDA library mode and library-free mode, but is worse when using the *in silico* predicted library. Spectronaut has the lowest precision of all workflows for the DDA library and library-free mode. The *in silico* predicted libraries does improve the precision of the analysis.

## Discussion

Although DIA tackles the problem of missing values, an inherent limitation of DDA which is biased towards more abundant peptides/proteins, it is not always clear how qualitative the extra data is. By spiking a dynamic range standard mixture of 48 human proteins (ranging 6 orders of magnitude) in different dilutions into a constant yeast background, the DIA data quality can be thoroughly evaluated. Nowadays, a plethora of options are available for DIA data analysis, each having an impact on the result outcome and quality. A total of 12 DIA workflows were evaluated and the performance compared to DDA and to each other. The workflows are combinations of the three commonly used DIA software packages Spectronaut, EncyclopeDIA and DIA-NN, each run with four different library methods (DDA-based spectral library, *in silico*-created libraries using the MS2PIP or the PROSIT algorithm, and library-free). The performance of the different workflows was evaluated on three criteria: detectability, precision and accuracy. In general, the detectability is substantially increased by using library-free mode (in DIA-NN) or when using *in silico* generated spectral libraries. EncyclopeDIA detects the UPS proteins fairly consistently among different dilutions, while DIA-NN suffers more from the dilution of the UPS proteins. DIA-NN however detects more background yeast proteins in all workflows. With EncyclopeDIA, the detectability is the best when using the PROSIT-predicted library, while DIA-NN equally performs in library-free mode as with the *in silico* predicted libraries. Although the detectability is superior with DIA for 11 out of the 12 workflows, the precision and accuracy is highly dependent on the workflow. More specifically, the lower abundant proteins that are not detected with DDA suffer more from higher variability, as can be expected because of their lower signal-to-noise ratio. DIA-NN shows a better precision in general, even outperforming DDA, independent of the workflow that is used. However, the accuracy of the DDA results on the UPS proteins outperforms most DIA workflows, despite the lower detectability. DIA-NN and EncyclopeDIA have comparable accuracy independent of the workflow and perform the best. Again, as is the case for the precision, the lower abundant proteins suffer more from low accuracy. The data suggests to be more careful with data from low abundant proteins and to apply a higher number of replicates in order to draw reliable conclusions. Nonetheless, DIA-NN in library-free mode or using *in silico* predicted libraries rendered the best result in this study, confirming the fact that DIA-NN is already optimized for *in silico* generated libraries (what it uses in its library free option). Considering the performance on the yeast proteome, DIA-NN in combination with an in silico predicted library or in library free mode performs the best, proving its success in combining both peptide centric and spectrum centric approaches.

Isaksson *et al.*^22^ state that the quality of the spectral library defines the quality of the quantification in DIA. Here we show that a DDA-created spectral library despite its higher quality, does not necessarily have a better precision nor accuracy. Despite the fact that *in silico* generated libraries increase the size of the spectral library substantially^22^, and hence suffer from a higher FDR, the quality is sufficient to outperform DDA libraries. Predicted libraries were shown here to increase the number of protein identifications, but because of the low abundancy of these extra identifications, the precision and accuracy is hampered. By further improving the quality of the *in silico* spectral library the identification and quantification of these proteins might be more reliable. For example, by adding a gas phase fractionation (GPF) analysis to filter the spectral library, the performance of this approach might be boosted because of a better FDR performance^37^. The same applies to the combined use of PROSIT, DIA-Umpire and DeepLC in the recently published MSLibrarian^22^. The quantification accuracy can be further improved as well by using fragment level ratio maximization and normalization^22^.

Different window sizes in the data acquisition may lead to different results^46^ because of the change in the number of points across the peak^24, 60^ on one hand and the higher chimeric content of the spectra on the other hand. As algorithms are still evolving, benchmarking new developments will remain necessary. Although for a long time it was considered that a minimum amount of points across the peak is needed, with 9 being the optimum according to Pino *et al.*^24^, Doellinger *et al.*^60^ recently stated that 1.5 points across the peak are sufficient and even increases the number of identifications. Our data showed 3 to 4 data points across the peak, depending on the workflow used. The resulting number of data points across the peak is highly depending on the chromatography as well^24, 60^. The application of DIA in clinical proteomics and single cell proteomics drives the technology towards shorter gradients in order to measure as many samples per day as possible. Such shorter gradients will have an effect on the performance of the different workflows. Hence for each type of application the performance should be checked with a same type of sample. And as software algorithms improve, a high chimeric content of spectra might be less of a problem in the future.

Post translational modifications (PTMs) are not taken into account in this data set. If the PTM does not interfere with the intensity ranking of the fragments, it should be possible to perform data analysis. In case the PTM does interfere with fragment intensity ranking, the spectral prediction algorithm needs to be trained first. The use of a DDA created library remains the safest option in the case of PTMs that alter the fragment intensity ranking, although with deep learning predicted spectral libraries phosphoproteomics is possible^61^

We can conclude that depending on the workflow, DIA renders more (low abundant) identifications as compared to DDA. A lower precision and especially lower accuracy inherent to the high number of low abundant identifications is inevitable though. The results obtained from this study apply to shotgun type of samples, and may differ when other types of samples are analyzed such as biological fluids or when the input is lower as is the case in single cell proteomics. Indeed, this benchmark resembles a shotgun sample present in abundant quantity. Jiang *et al.*^45^ showed in the case of low input samples that DIA-NN in combination with GFP-created libraries outperforms EncyclopeDIA and Spectronaut. Since fragmentation patterns can change in single cell analysis^62, 63^, depending on the detector and analyzer used^62^, another type of benchmark sample should be used. Although a faster analysis method should be used in the case of single cell proteomics since the sample numbers are large, it has been proven many times that DIA is capable of coping with fast analysis methods^60^.

## Supplementary figures

**Supplementary Figure 1.**
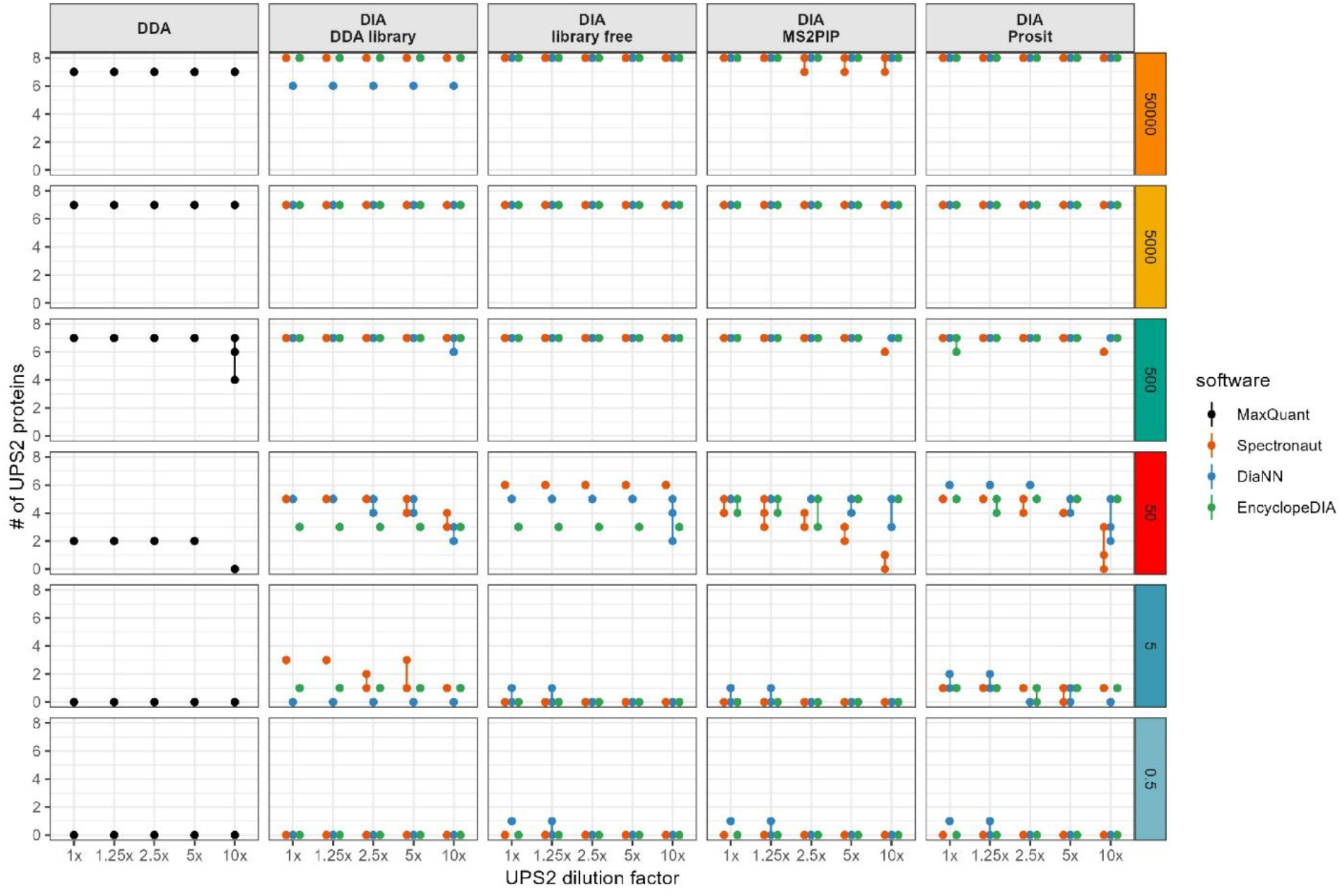
UPS2 proteins detectability according to analysis workflow, sample dilution and UPS2 protein abundance. Number of UPS2 mixture proteins identified in each analysis workflow, according to sample dilution in yeast extract (1x down to 10x) and protein abundance group (reference values “50000” down to “0.5”, representing groups of eight proteins at six orders of magnitude). Each software is represented with a different color Panel columns depict the three DIA library methods and the DDA workflow.

**Supplementary Figure 2.**
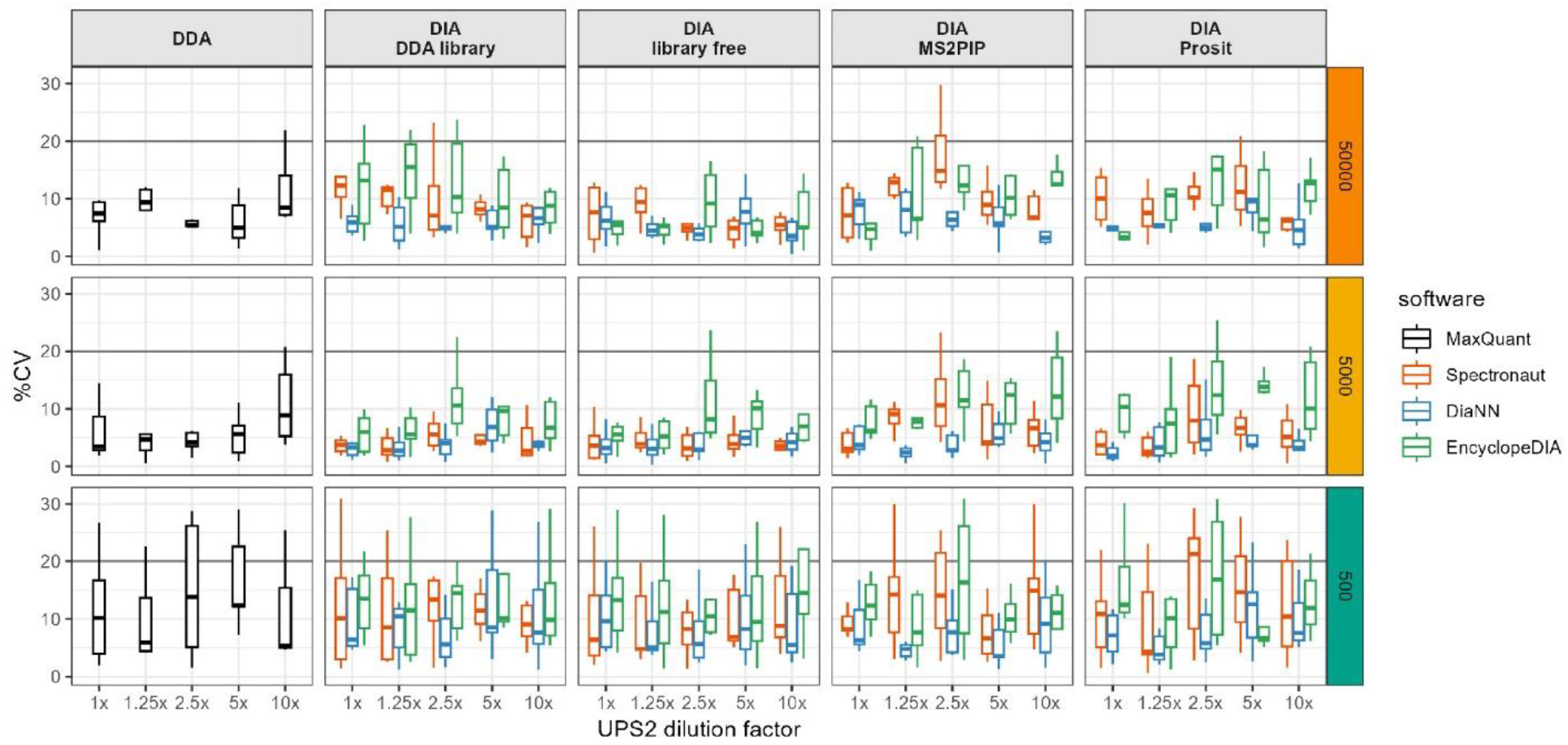
UPS2 proteins quantification precision according to analysis workflow, sample dilution and UPS2 protein abundance. Boxplots of percent coefficients of variation of UPS2 mixture proteins identified in each workflow of analysis, according to sample dilution in yeast extract (1x down to 10x) and protein abundance group (reference values “50000” down to “0.5”, representing groups of eight proteins at six orders of magnitude).

**Supplementary Figure 3.**
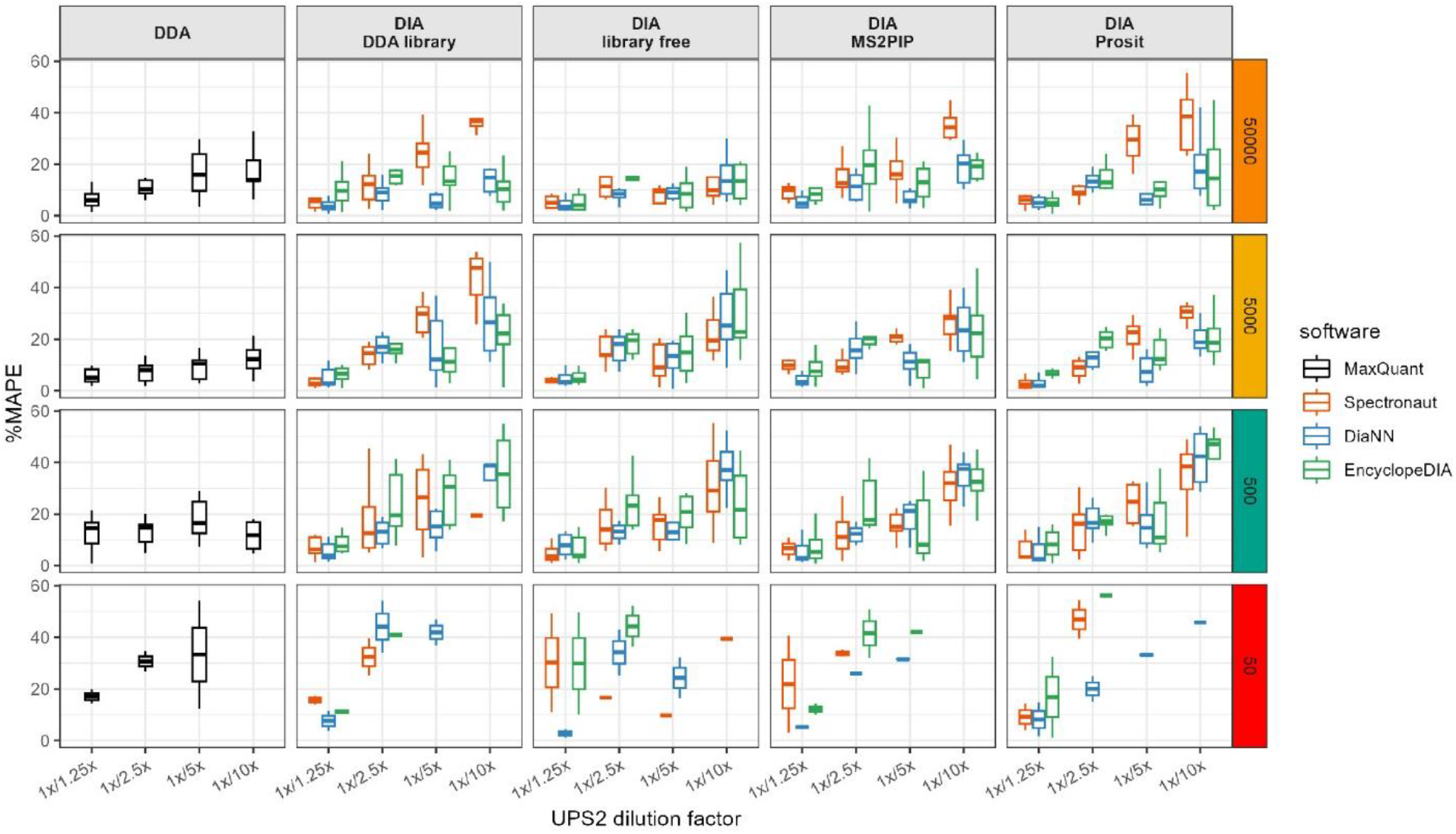
UPS2 proteins quantification accuracy according to analysis workflow, sample dilution and UPS2 protein abundance. Assessment of protein group quantification accuracy was done using the median absolute percentage error (MAPE) of the ratios of UPS2 protein intensities between the 1x yeast extract:UPS2 dilution relative to each of the other four sample dilutions (1.25x, 2.5x, 5x, 10x).

